# Non-invasive modulation of brain activity and behavior by transcranial radio frequency stimulation

**DOI:** 10.1101/2025.11.27.691064

**Authors:** Omid Yaghmazadeh, Leeor Alon, Tanzil M. Arefin, Zakia Ben Youss, Jiangyang Zhang, György Buzsáki

## Abstract

**Background:** Achieving non-invasive, targeted modulation of deep brain tissue remains a major challenge in neurotechnology. Current non-invasive brain stimulation methods—such as transcranial electrical (TES), magnetic (TMS), and focused ultrasound (TFUS) stimulation—suffer from limitations in spatial focality, penetration depth, or skull-related distortions. Radio frequency (RF) energy, which penetrates biological tissue effectively, offers an alternative avenue for neural modulation. This study introduces Transcranial Radio Frequency Stimulation (TRFS) as a novel, contactless neuromodulation technique that leverages RF-induced thermal effects to modulate neural activity in vivo.

**Methods:** We developed a custom RF stimulation system using 945 MHz stub antennas optimized for localized brain heating in mice. Using our unique experimental setup, we developed and tested two operational modes of TRFS:

<u>Pristine mode</u>: RF stimulation applied to intact brain tissue.

<u>RF-genetics mode</u>: RF stimulation applied to brain regions virally transduced to overexpress the thermosensitive TRPV1 ion channel.

Neural activity was recorded using metal-free one-photon fiber photometry with GCaMP calcium indicators. Behavioral effects were assessed through a rotational test in freely moving mice after MK-801-induced hyperlocomotion. Local temperature changes were monitored by optical thermometry.

**Results:** In pristine mode, RF exposure induced temperature rises leading to dose-dependent suppression of cortical parvalbumin (PV) interneuron activity. This neural suppression translated behaviorally into a unilateral rotational bias ipsilateral to the stimulated hemisphere in hyperlocomotive freely moving mice.

In RF-genetics mode, RF stimulation of TRPV1-overexpressing regions produced temperature-dependent excitation of neural activity once local change in temperatures exceeded ΔT ≈ 1.5 °C. Behaviorally, this excitation reversed the direction of rotation in hyperlocomotive freely moving mice, yielding a contralateral bias.

**Conclusions:** TRFS represents a conceptual advance in neuromodulation, uniting the inherent capability of RF energy to target deep brain tissue with the biophysical reliability of thermal modulation. TRFS applications are bimodal, capable of influencing the pristine brain by suppressing the activity of specific neuronal populations in targeted regions, or of exciting selectively transfected neural ensembles expressing thermosensitive TRPV1 ion channels. The latter modality, first introduced here, represents a novel concept termed “RF-genetics.” TRFS represents a promising platform for next-generation non-invasive brain stimulation with potential translational applications in treating various neurological and psychiatric disorders.

## Introduction

Brain disorders constitute a diverse and extensive array of conditions that impact a substantial portion of the global population, affecting approximately one in every six individuals worldwide^1^. A considerable fraction of patients afflicted by these neurological and psychiatric conditions develop resistance to pharmacological treatments in a few years. Non-invasive brain stimulation methods, when proven efficacious in treating such patients, hold paramount importance as a treatment alternative to chronic drug administration or surgical interventions.

Currently, non-invasive brain stimulation techniques, such as Transcranial Electric, Magnetic, or Focused Ultrasound Stimulation - TES, TMS, and TFUS, respectively, are widely employed in neuroscience research and clinical practice^2^. Nevertheless, despite the widespread adoption and utility of these established methodologies, each is associated with notable and inherent limitations. Specifically, the application of TES lacks precise spatial focality and is unable to effectively target deeper brain structures due to the significant shunting of electrical currents through superficial tissues^3^. Similarly, the applicability of TMS is largely restricted to cortical brain regions due to the rapid decay of magnetic field strength with increasing tissue depth^4^.

Furthermore, the efficacy of TFUS is often compromised by skull-induced acoustic heterogeneities, which can distort the ultrasound beam, and its stimulation volume is typically limited to relatively small regions^5^. Consequently, the ongoing pursuit of novel and innovative techniques in the field of neuromodulation is of considerable interest, as it provides a broader spectrum of potential non-invasive therapeutic applications for a wider range of brain disorders.

Since the initial applications of Radio Frequency (RF) energy in various scientific and medical contexts, the multifaceted effects of RF on biological tissues, encompassing both potential risks and beneficial health applications, have garnered significant and sustained interest from the scientific community^6–8^. The brain, along with the nervous system in general, due to its intrinsic electrical nature, has consistently been considered a particularly vulnerable organ to exposure to RF energy^9,10^. This interest has been further amplified by the pervasive development and widespread use of cellular phones, which are typically held in close proximity to the head, thereby increasing focus on RF’s potential effects on brain tissue. As a result, a substantial body of literature has been generated over several decades, documenting various effects of RF energy deposition on brain structure and function. In addition, and as a logical extension of these investigations, RF energy has been employed in several medical applications such as tissue ablation^11^, hyperthermia^12^, stroke detection^13^, cancer treatment^12^, among others^14^.

Despite this history of investigations and applications, the potential of RF energy for direct brain stimulation has not been thoroughly explored. Over the past few years, we have attempted to elucidate the potential effects of RF energy exposure on neural activity^15–17^, with the overarching goal of harnessing these effects for non-invasive brain stimulation. We first investigated whether non-thermal doses of continuous wave (CW) RF exposure could modulate brain activity. Our one-photon calcium (Ca^2+^) imaging experiments, conducted in awake, head-fixed mice using a custom-developed fiber-coupled, RF-interference-free head-mount endoscope^15^, demonstrated that RF exposures with strengths several times higher than the regulatory limits do not significantly affect neuronal activity^15^. While these findings suggest that such specific RF exposure parameters may not be suitable for direct neuromodulation purposes, the results of this initial study provide critical empirical evidence addressing a long-standing debate regarding the potential for non-thermal effects of RF energy exposure on neuronal function demonstrating the absence of such effects^15^.

It is well established that local temperature changes can modulate neural activity at single-cell and broader network scales^18,19^. A recent study examined the impact of localized temperature increases, induced by the overexposure of green light laser commonly employed in optogenetic stimulation protocols, on the ongoing activity of neurons from diverse cell types across various brain regions. The findings of this study indicated that, when it existed, the overall effect was predominantly a suppression of neural activity^18^. Localized temperature elevations have also been demonstrated to excite neural activity through the targeted local overexpression of the temperature-sensitive Transient Receptor Potential Vanilloid 1 (TRPV1) ion channel, a technique known as magneto-genetic stimulation. Recent investigations have shown that localized heating of brain regions previously transduced with viral vectors for TRPV1 overexpression and subsequently loaded with ferromagnetic nanoparticles, also delivered via viral vectors, can lead to robust neural excitation upon the application of an external magnetic field^20,21^. Concurrently, it is well-documented that exposure to RF energy can effectively induce a measurable increase in the temperature of biological tissues^22^. Combining these concepts, we hypothesize that transcranial RF energy exposure, when applied to induce controlled temperature rises within the brain tissue, can be employed as a contactless neuromodulation technique capable of inducing both suppression and excitation of neural activity. In this study, we present proof-of-concept evidence supporting these possibilities through a series of in vivo experiments conducted in awake, behaving mice. We apply TRFS in two distinct modes: 1) the ‘Pristine mode’ in which TRFS is applied to the intact brain, and 2) the ‘RF-genetics mode’ in which TRFS is applied to the brain with an overexpression of TRPV1 in the target area.

Our previous experiments have shown that using metallic electrodes for electrophysiological recordings of neural activity can lead to falsified results, even when no visible artifacts are present^15^. Therefore, to achieve accurate results, it is crucial to employ metal-free recording methods for studying the effects of RF energy exposure on neural activity^15^. To assess neural activity during the experiments described in this report, we initially employed the blood-oxygen-level-dependent (BOLD) signal obtained from functional Magnetic Resonance Imaging (fMRI), with the primary aim of achieving a whole-brain readout of neural activity. However, as will be elucidated in this paper, the fMRI BOLD signal proved to be a less-than-ideal measure, particularly in the context of temperature alterations within the brain tissue. Consequently, we transitioned to optical calcium (Ca^2+^) imaging approaches. We initially attempted to use a fiber-coupled head-mount endoscope system that we had previously developed, based on the UCLA Miniscope V3 system, specifically designed for RF-interference-free electrophysiological recording^15^. However, as we were trying to induce temperature rises on a relatively fast time-scale, we had to use higher power of RF energy radiation compared to our previously reported experiments^15^. These elevated power levels caused the complete shutdown of the Complementary Metal-Oxide-Semiconductor (CMOS) camera integrated into the head-mount endoscope system, thereby interrupting the ongoing recordings. To circumvent this technical limitation, we ultimately adopted 1-photon fiber photometry recording. This method avoids the presence of any electronic components near the RF stimulation source. While lacking single-cell resolution, fiber photometry provides a sufficiently sensitive and robust measure of neural population activity adequate for the specific aims and experimental parameters of the current study.

## Results

We utilized RF energy exposure to modulate ongoing neuronal activity in a manner that can be leveraged for potential therapeutic neuromodulation applications. Learning from our previous experiments^15^, any assessment of the effects of RF energy exposure on brain activity should be performed in metal-free preparations.

### RF stimulation of the mouse brain

In our previous study, we used a Transverse ElectroMagnetic (TEM) cell and a Patch antenna which provided full body exposure ^22^. Because in the present study, we aimed to induce local temperature increase, we designed an antenna, stub antenna Version 1 (V1), composed of an open-end coaxial RF cable with an unshielded tip (of ∼1cm length) which is matched by a simple printed circuit board (PCB) at 945MHz (all the experiments in this study are performed at this frequency) including a transmission line and a modifiable parallel capacitor trace (Fig. 1A). This antenna was designed for experiments with head-fixed animal in which it was positioned on the animal’s head and kept in place mechanically (e.g., using a headcap with a designated position for the antenna tip). We first tested whether the stub antenna induces temperature rises in the animal’s brain but not in its body (Fig. 1B; the mouse was anesthetized to avoid discomfort). Fig. 1C shows the measured temperature from the brain, abdomen and rectum when five trials of 15s RF_ON, followed by 81s RF_OFF recovery periods were applied. In sum, RF stimulation by the stub antenna induced selective heating of the brain (Fig. 1D).

**Figure 1.**
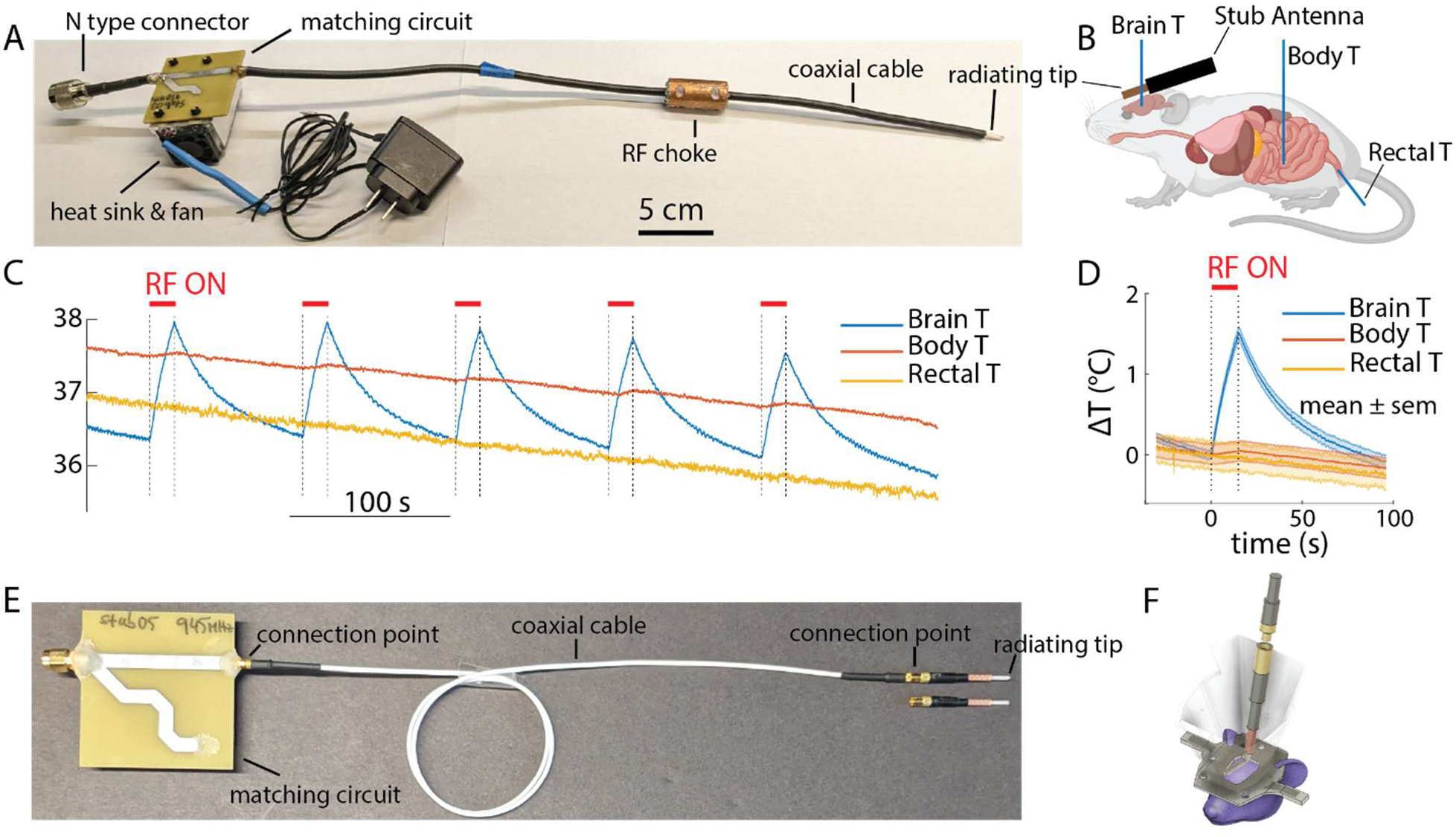
Stub antenna for selective RF stimulation of mouse brain. **A)** Photograph of the custom-made stub RF antenna (v1) for local stimulation of the mouse brain. **B)** An illustration of the antenna’s placement in a head-fixed mouse set-up. **C)** The temperature measurements from the brain (Brain T), the abdomen (Body T), and the rectum (Rectal T) under RF stimulation (trials of 15s RF-ON followed by 81s RF-OFF) using the custom-made stub antenna. **D)** Averaged temperature profiles (mean ± standard error of the mean (sem) across all RF stimulation trials. **E)** Photograph of the stub RF antenna (V2), which uses a mini-RF-connector so that the front end can be fixed to animal’s head for consistent stimulation. **F)** Schematics of animal’s head with an antenna V2 attached. With V2, RF stimulation can be performed in head-fixed or freely moving mice.

To make the antenna suitable for chronic experiments in waking mice, we improved the antenna design (stub antenna Version 2 (V2)): a short antenna tip is attached to animal’s skull and the rest of the circuitry (Fig. 1E). When stimulation is not applied, the coaxial cable can be disconnected from the tip (which is protected in a 3D-print head cap also attached to animal’s skull; Fig. 1F). Fixing the antenna tip to the skull resulted in consistent and repeatable stimulation patterns over different experiments. In addition, using a lighter coaxal cable enabled us to perform experiments with freely moving animals. We used these set-ups in the resto of the experiments to study the thermally mediated effects of RF energy exposure on neural activity.

### Magnetic Resonance Imaging (MRI) assessment of the effects of TRFS in vivo

In our initial approach to assess neural activity under RF stimulation, we employed simultaneous functional Magnetic Resonance Imaging (fMRI) and RF stimulation in anesthetized mice (Suppl. Figs. 1 and 2). In summary, we observed both increased and decreased Blood-Oxygen-Level Dependent (BOLD) signals in various brain regions (Suppl. Fig.1). However, these experiments were hard to interpret because BOLD decrease was possibly due to changes in the temperature-dependent MRI parameters, such as T_2_, and local blood flow in the brain. Therefore, although these results showed TRFS-induced physiological changes in the brain, we were unable to extract net neural effects from the BOLS signal. To achieve a more precise evaluation of TRFS effects on neuronal activity, we next employed optical Ca^2+^-imaging.

**Figure 2.**
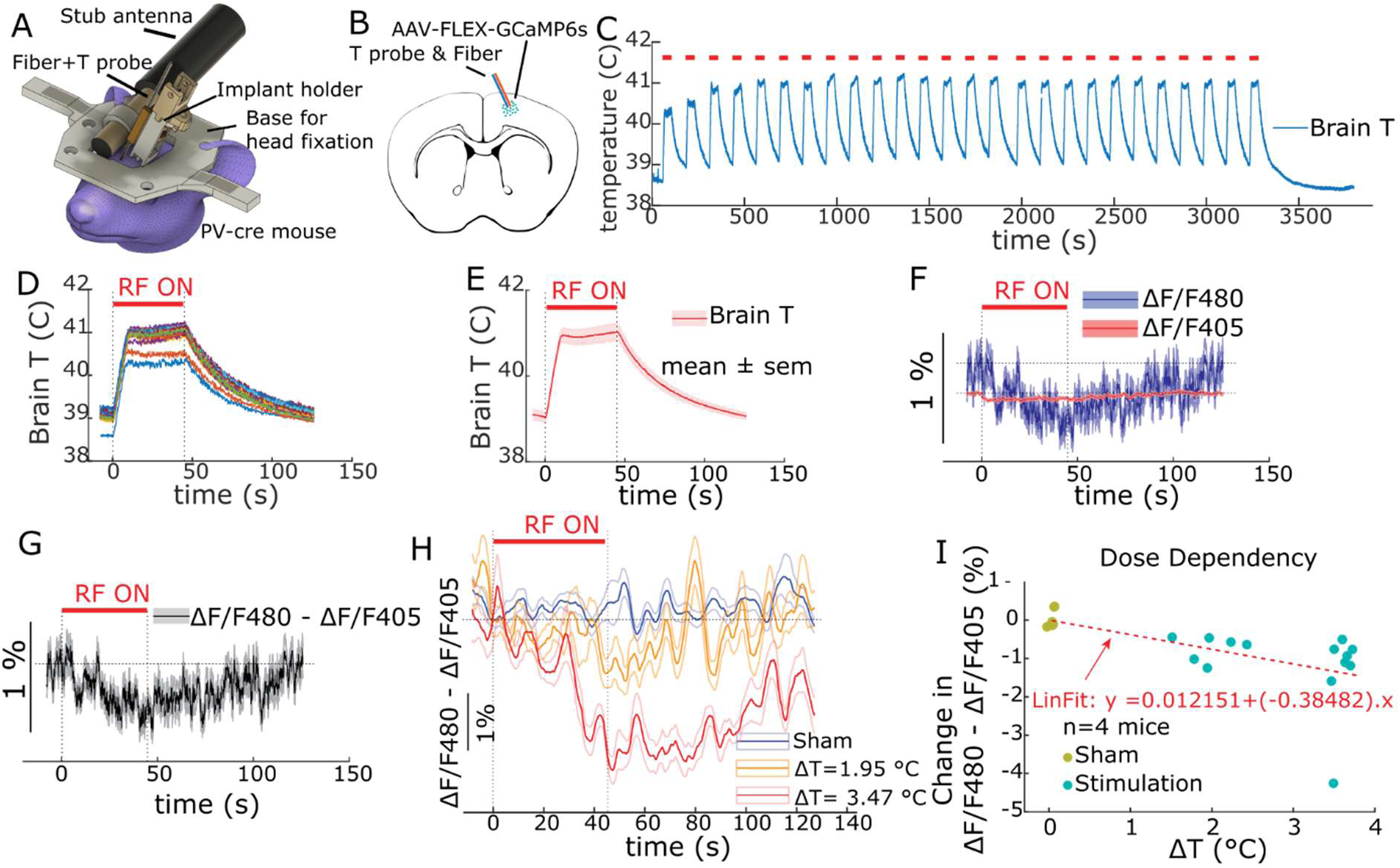
TRFS-induced suppression of neural activity in *pristine mode*. **A)** Graphical presentation of the animals’ head after head-cap attachment, fiber (for fiber photometry) and optical temperature sensor implanted, and the RF stub antenna positioned in place for stimulation. **B)** Illustration showing the mouse brain slice with the virus injection site. Note that PV-cre mice were used. **C)** Temperature profile of an example recording session where trials of 45s RF-ON followed by 81s RF-OFF periods were applied. **D)** Overlaid temperature profiles of the trails from C. **E)** Average (mean ± sem) temperature profile of the trials in C. Different colors show data from different trials. **F)** Average fiber photometry response (mean ± sem) for all trials including the main excitation signal (480 nm, blue) and the isosbestic control signal (405 nm, red). **G)** Average fiber photometry response (mean ± sem) with isosbestic correction. **H)** Ca^2+^ responses at different strengths of the RF stimulation. Inset indicated the induced temperature rises. **I)** Correlation between the amplitude of the fiber photometry response and the RF-induced temperature-rise (with a linear fit curve) in n=4 PV-cre mice. Each dot corresponds to a recording session.

### TRFS-induced changes of neural activity in the intact brain in vivo

Guided by recent reports in the literature on temperature-rise-induced suppression of neural activity^18^, we examined whether using the thermal effect of TRFS we could induce non-invasive neuromodulation (n=4 mice; Fig. 2). We examined such temperature-driven effects on cortical fast spiking PV interneurons (using a PV-cre mouse line), because PV interneurons have been shown to have a to have pronounced response to temperature rise^18^. Fig. 2C-E summarizes the temperature changes in the brain in response to RF stimulation. Simultaneously measured fiber photometry signals, reflecting summed activity of PV interneurons are shown in Fig. 2F, G. RF-induced heating decreased the activity of cortical pyramidal PV interneurons in a dose-response manner (Fig. 3H, I).

**Figure 3.**
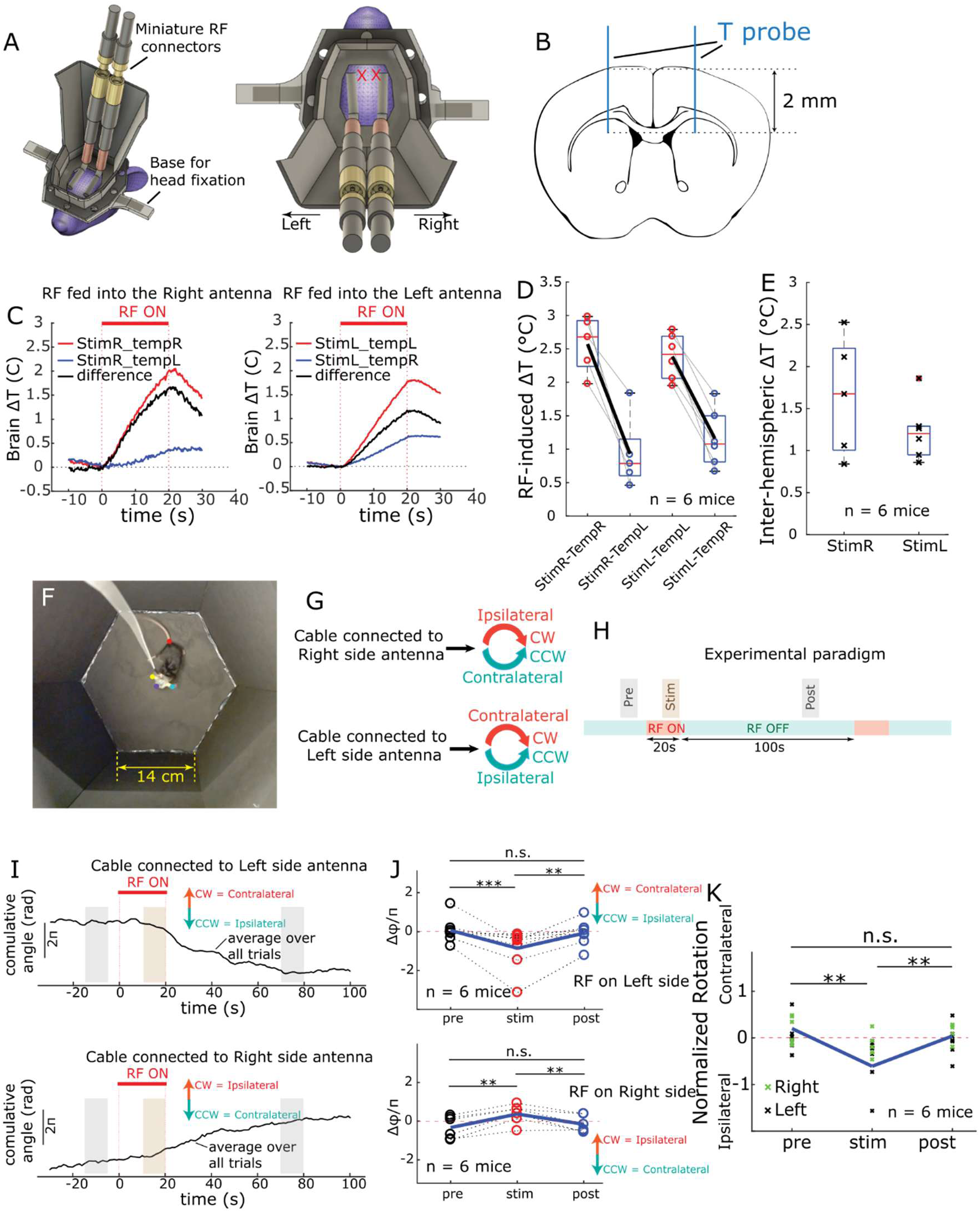
TRFS-induced behavioral changes via suppression of neural activity in *pristine mode*. **A)** Illustration of example mouse head with attached stub antennas on top of both brain hemispheres. Red crosses on the right plot show the location where temperature measurements are performed. **B)** A representative mouse brain slice indicating the location of the optical probes to measure induced temperature rises at striatal and cortical depths. **C)** Temperature measurement results showing a differential heating of striatal regions of each hemisphere due to right or left side RF stimulation (example data from one animal). **D)** Temperature rises measured from the striatum region (indicated in B) from both right and left hemispheres when the RF stimulation is applied on the right or left side for all animals (n = 6 mice, 3 male and 3 female). **E)** RF-induced hemispheral temperature difference as a result of RF stimulation applied to the left or right side (same data as D). **F)** Example frame from recorded video showing the open-field rotation behavior experiment. Small colored circles indicate computationally detected animal’s body parts used for data analysis. **G)** illustration that indicates the direction of ipsi- or contralateral movements depending on which antenna (right or left side) is used for stimulation. **H)** Experimental trials design consisting of 20s RF-On followed by 100s RF-OFF; intervals with indication of ‘pre’, ‘stim’, and ‘post’ periods used for comparative analysis. **I)** Example traces of animal’s angular displacement (measured by calculated cumulative angle of animal’s head) averaged over all trials in sample sessions where RF was applied either to the left (top) or the right (bottom) antenna. On each plot, arrows show the anticipated change in cumulative angle for CW and CCW rotations. **J)** average change in cumulative angle per trail for all animals for pre, stim, and post periods for both left (top) and right (bottom) side RF stimulation cases (n=6 mice, 3 male and 3 female). **K)** Normalized rotational direction for all animals and combining the data in the two plots in J (n=6 mice, 3 male and 3 female).

### TRFS-induced behavioral changes via suppression of neural activity in the intact brain

A compelling way to evaluate the effectiveness of any neuromodulation technique, such as TRFS, is to demonstrate its capability in altering the animals’ behavior. To achieve this goal, we employed a rotation test in freely moving mice, following a previously established protocol^15^. Mice were placed in a circular open-field environment shortly after being injected with MK-801 (0.30 mg/Kg weight). MK-801 (a.k.a. Dizocilpine) is a non-competitive antagonist of the N-methyl-D-aspartate (NMDA) receptor, and its injection induces hyper-locomotion in mice^23^. As a result, mice ran in circular patterns, in both clockwise (CW) and counterclockwise (CCW) directions, for a substantial period of time (∼ 1 hour)^23^. Our experimental design was based on previous experiments, in which manipulating the temperature unilaterally in the dorsal striatum induced suppression of neural activity in medium spiny neurons (MSNs), resulting in a preference for rotational behavior ipsilateral to the manipulated hemisphere^18^. We attached two stub antenna V2s to the mouse skull to target left and right hemispheres separately (Fig. 3A, B). Feeding one antenna at a time allowed unilateral RF stimulation of the brain (20s followed by a period of at least 100s of no stimulation). To ascertain the efficacy of RF stimulation, we first measured the induced temperature changes in both hemispheres when RF was applied unilaterally. For these measurements, mice were head-fixed, and temperature probes were temporarily inserted into their brain through formerly prepared craniotomies at AP: +1mm and ML: ±1.5mm coordinates from the bregma point (n = 6 mice; Fig. 3B-E). While unilateral RF stimulation increased temperature in both hemispheres of the brain, the difference exceeded 1°C (Fig. 3D-E). We hypothesized that the resulting imbalance of neuronal activity changes (as shown in Fig. 2) should lead to measurable behavioral effects.

To verify this hypothesis, we tested the implanted mice in an open-field arena (Fig. 3F). Locomotion was monitored by video recording (30 frames per second) and directional changes were calculated from the animal’s body parts locations, as determined using Deeplabcut^24^. Fig. 3G shows the corresponding ipsi- or contralateral rotations, in relation to the stimulated hemisphere. The experimental trial design is shown in Fig. 3H: in each session, RF is unilaterally applied in several (20-30) trials of 20s ON followed by 100s OFF periods. Measured data were quantified and compared between 10s long ‘Pre’, ‘Stim’ and ‘Post’ periods. The cumulative angle of the animal’s head varied, showing a rotational direction preference ipsilateral to the side of RF stimulation (n=3 male and 3 female mice. Fig. 3I-K; see Suppl. Vid. 1 and Suppl. Vid. 2, for example videos. These results demonstrate that TRFS *pristine mode* can effectively alter the rotational behavior of the mice.

### TRFS-induced excitation of neural activity in vivo via TRPV1 overexpression in RF-genetics mode

We observed a reduction of neuronal activity in the neocortex and demonstrated behavioral changes by application of TRFS on the intact brain. Next, we hypothesized that TRFS can also be used to excite neurons based on previous observations that heat can increase spiking activity in neurons via overexpression of temperature-sensitive ion channels (such as TRPV1) ^20,21^. We injected an adeno-associated virus (AAV) construct (AAV9-SYN-mTRPV1-P2A-mCherry) into the motor cortex and used RF-artifact-free fiber photometry to assess the ongoing neural activity (n=4 wild mice; Fig. 4A-B). The RF stimulation (20s RF-ON followed by 100s RF-OFF periods) raised the brain temperature (Fig. 4C,D) and lead to an increase in neural activity, as shown be elevated GCaMP excitation (Fig. 4E,F). The magnitude of the fiberphotometric response varied with the applied RF power and the induced brain temperature (Fig. 4G). Fig. 4H and 4I show the relationship between the amplitude of the neural response to RF stimulation and the corresponding induced temperature rise (ΔT) and the maximum attained temperature (T_max_), respectively. The data distribution reflects a linear relationship between the response and the respective temperature parameters (ΔT or T_max_) above a threshold value. To estimate these threshold values, we used linear regression on data in its linear regime (higher values of ΔT or T_max_) and calculated the x-intercept of the fitted line. This analysis indicates that threshold values of 1.53 **°**C and 39.37 **°**C, for ΔT and T_max_, respectively, are needed to obtain a neural response.

**Figure 4.**
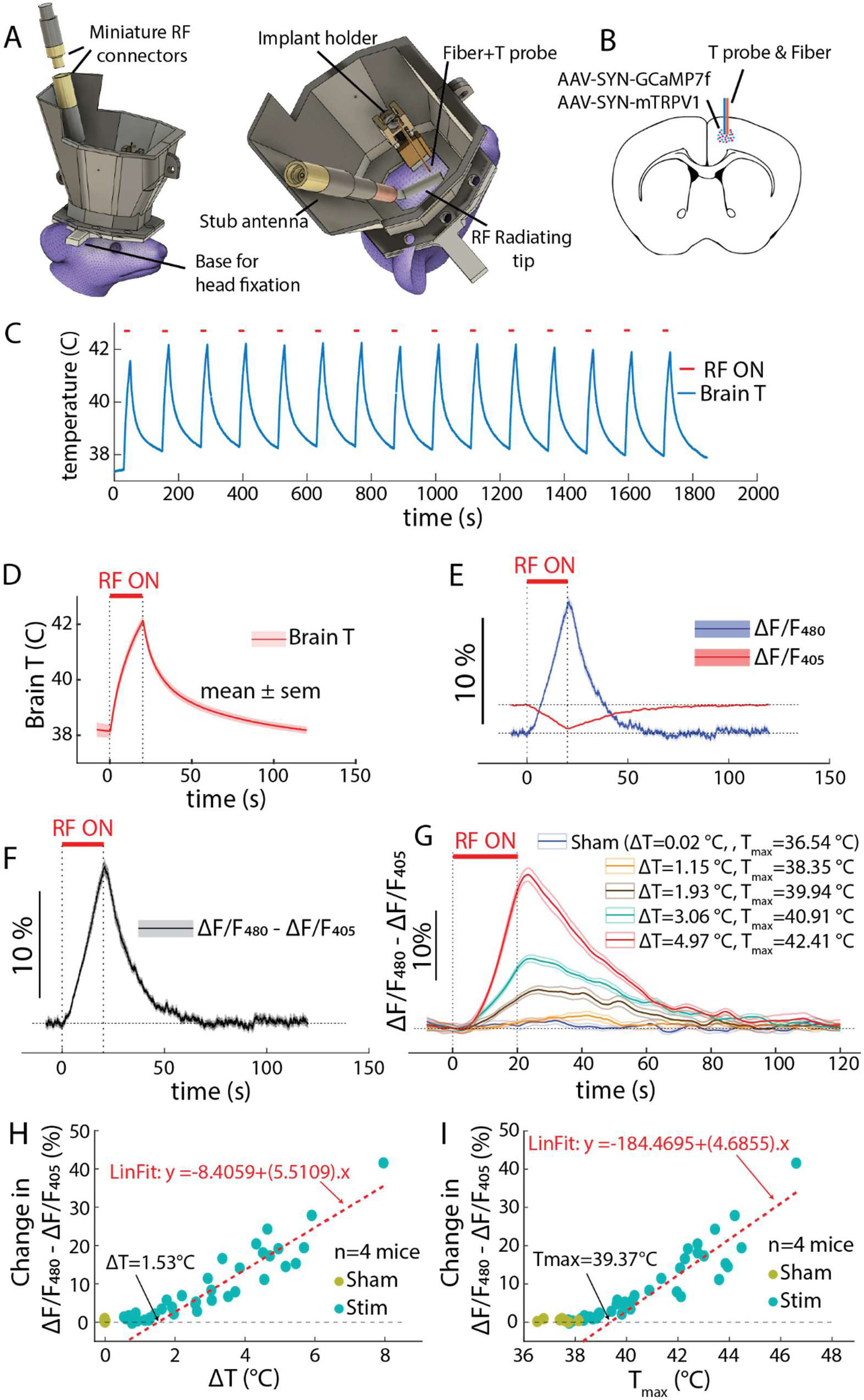
TRFS-induced excitation of neural activity via overexpression of TRPV1 channels in *RF-genetics mode*. **A)** Graphical presentation of the animals’ head after head-cap attachment, fiber (for fiber photometry) and optical temperature sensor implanted, and the front part of the RF stub antenna for stimulation. **B)** Illustration showing the mouse brain slice with the virus injection and implantation site. **C)** Temperature profile of an example recording sessions where trials of 20s RF-ON followed by 100s RF-OFF periods are applied. **D)** Average (mean ± sem) temperature profile over all the trials in C. **E)** Average fiber photometry response (mean ± sem) for all trials including the main excitation signal (480 nm, blue) and the isosbestic control signal (405 nm, red). **F)** Average fiber photometry response (mean ± sem) with isosbestic correction. **G)** Dependency of the neural response to the strength of the RF stimulus and the resulting temperature rise (ΔT) and the highest temperature (T_max_) in recordings from one mouse. **H, I)** Relationship between the amplitude of the fiber photometry response and the RF-induced temperature-rise (ΔT) and the highest reached temperature (T_max_) in n=4 wild-type mice.

We also examined persistence of TRPV1 expression and the efficacy of the neurostimulation effects over time. We repeated the experiment 7 months after the virus injection and obtained similar results compared to measurements performed 4 weeks after the injection (the waiting time for viral expression; n=1 mouse, Suppl Fig. 3), illustrating the stability of the stimulation effects.

### TRFS-induced behavioral changes via excitation of neural activity in RF-genetics mode

Similar to the TRFS *pristine mode* in the intact brain, we employed the same rotational behavior test to examine the efficacy of TRFS *RF-genetics mode*. In a new cohort of the animals, we virally overexpressed TRPV1 channels in the striatum unilaterally (n= 3 male and 3 female mice; Fig. 5A, B). All other aspects of the experiments were identical to those of the behavioral test in *pristine mode* (Fig. 3). Similar to experiments conducted in the brains of animals lacking TRPV1 channel expression, hemisphere-specific temperature rises were observed in the striatum (Fig. 5C, D). In the open-field test, RF was applied in 20 to 30 trials of 20s ON epochs followed by 100s OFF epochs. Ten-second-long ‘Pre’, ‘Stim’ and ‘Post’ periods were used for data comparisons (Fig. 5E, F). The results showed opposite rotational changes (Fig. 5G, H; Suppl. Vid. 3 and Suppl. Vid. 4) than in animals without TRPV1 channel expression (compare with Fig. 3). As an additional control, we show that when RF stimulation was delivered to the hemisphere without TRPV1 channel expression, the animals rotated toward the RF-stimulated side, mirroring the rotation behavior results in non-injected mice (Fig 5H in comparison with Fig. 3).

**Figure 5.**
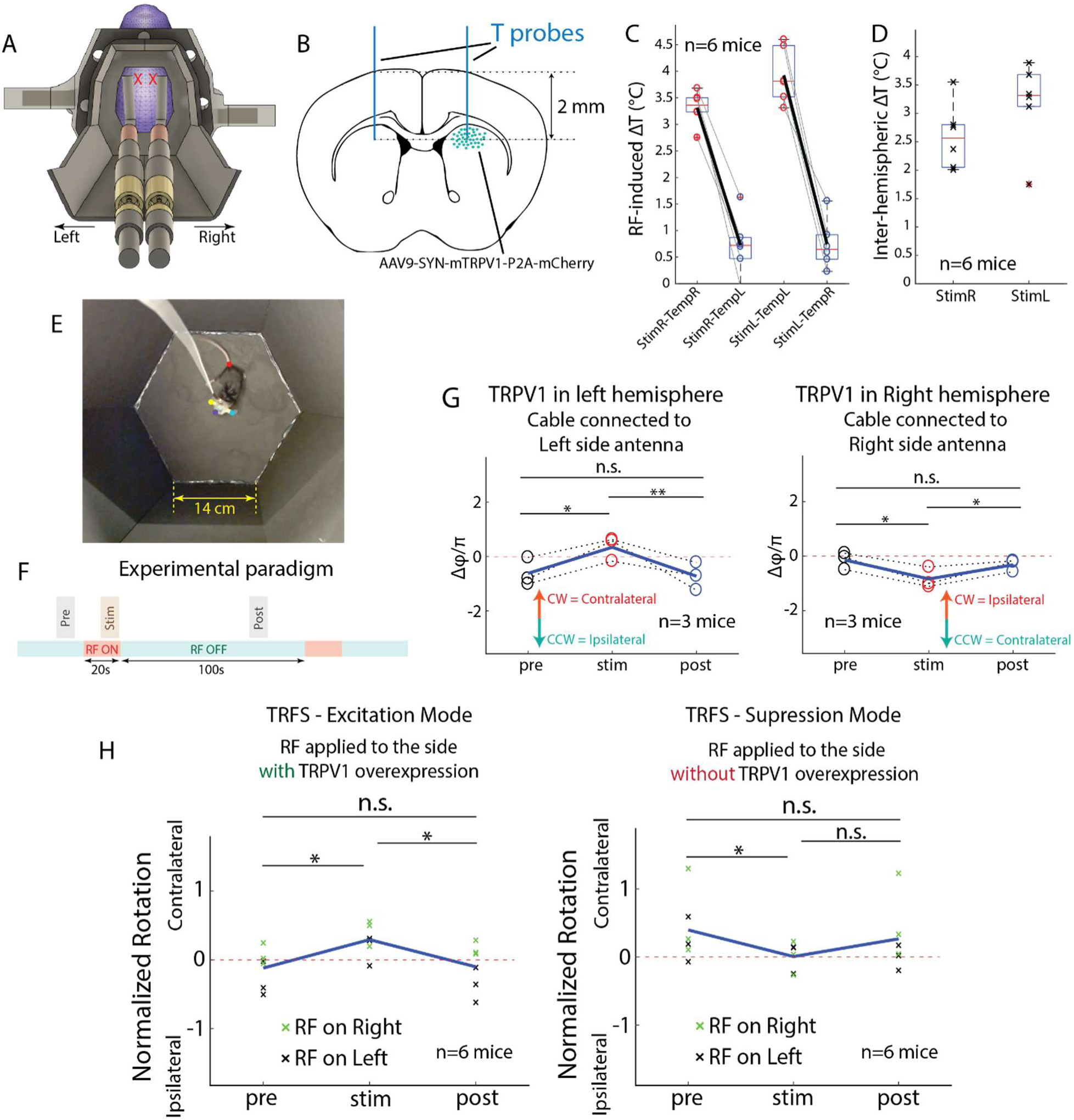
TRFS-induced behavioral changes via excitation of neural activity in *RF-genetics mode*. **A)** Schematics of bilaterally implanted RF antennas and head stage. X, location of temperature measurements. **B)** Mice were injected unilaterally (n=3 in the left and n=3 in the right or left hemisphere) in the striatum to induce overexpression of TRPV1 ion channel. **C)** Temperature rises measured in the striatum region when the RF stimulation is applied above the ipsi- or contralateral hemisphere (n = 6 mice, 3 male and 3 female). **D)** RF-induced hemisphere-specific temperature increase. **E)** Example frame from recorded video showing the open-field rotation behavior experiment. Small colored circles indicate computationally detected animal’s body parts used for data analysis. **F)** Experimental trials design consisting of 20s RF-On followed by 100s RF-OFF intervals with indication of ‘pre’, ‘stim’, and ‘post’ periods used for comparative analysis. **G)** Average change in cumulative angle per trial for pre, stim, and post periods for TRPV1 expressions in the left (left plot, n=3 mice) and the right (right plot, n=3 mice) hemisphere with RF applied to the same side. **H)** Normalized rotational direction for all animals and combining the data when stimulation is applied to left and right side for the following cases: 1) RF applied to the side with TRPV1 overexpression (TRFS *RF-genetics mode*; left plot, n=6 mice, 3 male and 3 female) and 2) RF applied to the side without TRPV1 expression (TRFS *pristine mode*; right plot, n=6 mice, 3 male and 3 female).

## Discussion

We introduce a novel methodology for achieving non-invasive neuromodulation through the controlled application of radiofrequency (RF) energy exposure. We established a proof-of-concept for achieving both suppression and excitation of neural activity within the brain through TRFS-induced controlled thermal manipulations. We demonstrated these neuromodulatory effects through *in vivo* neural activity measurements using fiber photometry in awake, head-fixed mice, as well as through a rotational behavioral assay performed in freely moving mice exhibiting drug-induced hyperlocomotion. Based on these findings, we delineate two distinct operational modalities for TRFS: the *pristine mode*, wherein localized temperature elevations induced by RF energy can effectively reduce the ongoing neural activity in some neuronal populations; and the *RF-genetics mode*, in which similar RF-induced temperature increase can selectively elicit neuroexcitation in neuronal populations that have been previously transduced with neuron-specific viral vectors to achieve overexpression of TRPV1 channel. The term *RF-genetics mode* is chosen to draw a clear parallel with other neuromodulatory techniques that leverage genetic modifications in conjunction with specific stimuli, such as opto-genetics (light), magneto-genetics (magnetic fields), and sono-genetics (ultrasound).

A particularly compelling feature of the *RF-genetics mode* of TRFS is its inherent potential for achieving cell type-specific targeting, a capability that is primarily driven by the sophisticated genetic tools employed in viral vector-mediated gene expression, as has been demonstrated in other genetically mediated neuromodulatory approaches^25^. We used fiber photometry to demonstrate the effectiveness of TRFS *RF-genetics* mode in exciting neural activity in cortical neurons in awake mice (Fig. 4). In contrast, the TRFS *pristine mode*, while effective in modulating neural activity through localized heating, appears to exhibit a more limited degree of neuron-type and brain-region specificity, as evidenced by existing literature detailing the differential effects of temperature changes on various neuronal populations. Notably, a recent investigation conducted by Owen and coauthors^18^ examined responses of neural activity to temperature increases induced by light stimulation across diverse brain regions in mice. Their findings revealed that the activity of cortical interneurons, striatal medium spiny neurons (MSNs), and dentate gyrus granule cells was significantly attenuated by temperature increase, whereas the activity of cortical pyramidal cells exhibited a slight decrease, and the activity of hippocampal CA1 pyramidal cells remained largely unaffected. Our fiber photometry results, demonstrating a significant *reduction* in neural activity of PV interneurons during TRFS-induced heating (Fig. 2), are consistent with these prior observations. Nevertheless, despite this inherent characteristic of the TRFS *pristine mode*, it remains a valuable tool for brain stimulation due to its ability to alter the activity of interneurons, which play a crucial role in maintaining the delicate excitatory-inhibitory (E/I) balance within neural circuits and in orchestrating overall brain function. In fact, a growing body of evidence indicates that the targeted suppression of specific interneuron populations can yield beneficial therapeutic impacts in the treatment of a range of complex brain disorders, including schizophrenia^26^, depression^27^, anxiety disorders^28^, and potentially other neuropsychiatric conditions. This understanding underscores that, despite its specific limitations targeting neuronal populations, the TRFS *pristine mode* represents a potent and impactful tool for non-invasive brain stimulation interventions. The mechanism of the neuromodulatory effects induced by *RF-genetics mode* is via thermal activation of the locally transduced temperature-sensitive TRPV1 channel, as also shown in magneto-genetics approaches^20^. For the effects induced by the *pristine mode*, a plausible mechanism, previously demonstrated by Owen et al.^18^, is via thermal activation of thermosensitive inwardly rectifying potassium (Kir) channels.

Beyond assessing TRFS’s neuromodulatory effects via fiber photometry, we also demonstrated its capacity, in both modes, to alter animal behavior through brain stimulation. We utilized a rotational assay in freely moving, drug-induced hyperlocomotive mice, leveraging the principle that unilateral stimulation of striatal neurons can bias directional preference. This behavioral paradigm has previously been employed by Owen et al.^18^ to demonstrate neuromodulatory effects of heating the striatum brain region. Their study showed that unilateral striatal light exposure, inducing a differential temperature increase, suppressed MSN neurons, which constitute approximately 90% neurons in this region^29^, leading to an ipsilateral directional preference relative to the stimulated side. We adopted a similar strategy. Upon unilateral TRFS application in its *pristine mode* to a striatal region (either left or right hemisphere), we similarly observed a rotational preference ipsilateral to the stimulation side (Fig. 3K). This finding aligns with Owen et al.’s observation that striatal heating induces suppression of activity in MSN neurons and an ipsilateral directional preference^18^. Conversely, when we applied TRFS *RFgenetics mode*, we observed a rotational preference contralateral to the stimulation hemisphere, opposing the effects observed in TRFS *pristine mode* (Fig. 5H). This outcome was expected, as the TRFS *RFgenetics mode* is designed to excite neuronal activity in the target region. A potential concern is whether the observed effects during TRFS *pristine mode* might arise from heating-induced changes in the cortical regions overlying the striatum, rather than from striatal stimulation itself, particularly given that RF-induced heating in our preparation was not entirely region-specific. However, this concern is mitigated because similar cortical areas are heated in experiments employing the TRFS *RFgenetics mode*, where stimulation is specifically targeted to the striatum via TRPV1 channel expression. If cortical effects were solely responsible for the ipsilateral rotational preference in the TRFS *pristine mode*, then their similar presence during TRFS RFgenetics mode application would either induce a confounding ipsilateral bias or attenuate the expected contralateral preference, which was not observed.

One fundamental objective of biomedical research is to translate innovative technologies into practical human applications that improve health outcomes. In alignment with this aim, our ultimate goal is to translate the promising neuromodulatory effects of TRFS into effective therapeutic interventions for neurological and psychiatric disorders. Achieving this objective requires the development of a sophisticated system capable of delivering targeted RF stimulation within the human brain. Notably, RF energy in the microwave range has a well-established history in inducing localized or whole-body hyperthermia for cancer^14,30^. Recent advancements, such as the development of a 72-dipole antenna array system for targeted human brain hyperthermia^31^, provide a valuable foundational framework for a dedicated human TRFS system. Furthermore, owing to their small and flexible form factor, an array of stub antennas offers the potential for designing smaller, lighter, and more comfortable wearable devices, thereby facilitating prolonged therapeutic applications.

The degree of invasiveness, primarily defined by the necessity for surgical intervention, is a critical factor for the translational applicability of any neuromodulation technique. While the *in vivo* experiments presented here required surgical procedures for recordings and targeted TRPV1 expression, human TRFS applications inherently offer a distinct non-invasive advantage. A human TRFS system can be engineered for MRI compatibility, enabling non-invasive validation of brain tissue targeting via MR thermometry, as demonstrated in prior RF-based hyperthermia work^31^. Furthermore, for the TRFS *RF-genetics mode*, non-invasive drug and gene delivery methods, e.g. by using focused ultrasound^32,33^, can ensure the entirely non-invasive nature of the therapeutic process. In contrast to other emerging electromagnetic field-based neuromodulation approaches (e.g., magnetogenetics), which rely on the local delivery of synthetic nanoparticles, TRFS’s *RF-genetics mode* utilizes only the overexpression of endogenous ion channels, such as TRPV1. This distinction significantly mitigates concerns regarding unknown long-term side effects associated with exogenous materials. While viral vectors remain a delivery method for genetic approaches, the growing confidence in their safety and efficacy for treating brain disorders^34^ further supports TRFS’s substantial translational potential and eventual clinical acceptance as a novel non-invasive neuromodulation strategy.

A notable feature of TRFS is its bimodality, which enables the modulation of neuronal activity in two distinct operational modes: *pristine* and *RF-genetics*. This capacity for bimodal control is paramount for treating the diverse and heterogeneous landscape of brain disorders, as effective therapeutic interventions often necessitate either the enhancement or attenuation of activity in specific target neuronal populations. For instance, targeted excitation of the subthalamic nucleus (STN) consistently improves motor symptoms in Parkinson’s disease^35^, while suppression of the activity of interneurons in the mPFC region can alleviate depressive symptoms^27^. Building upon these examples, TRFS holds substantial promise as a novel therapeutic modality for various neurological and psychiatric conditions. Another important feature distinguishing TRFS is its potentially extensive range of spatial coverage within the brain when used in human applications. As previously demonstrated by modeling and in phantom preparations, RF energy can be precisely focused by an antenna array to target a spherical volume of approximately 2 cm in diameter within deep human brain tissue^31^, capable of inducing a sufficient temperature rise for neuromodulation (2 °C), which is consistent with our experimental results. Furthermore, simultaneously driving multiple antennas within an array makes it feasible to influence significantly larger volumes, potentially encompassing the entire brain. This broad spatial span of influence positions TRFS as a unique non-invasive brain stimulation method, offering an unprecedented degree of flexibility for diverse targeting strategies—from focal deep brain regions to more diffuse, larger brain volumes—which can be tailored to specific neurological circuits and pathological conditions.

Current non-invasive brain stimulation methods, while offering a range of beneficial features for modulating neural activity, are each inherently constrained by distinct limitations arising from the fundamental nature of their respective stimuli and their interaction with biological tissues. Consequently, developing novel NIBS methods remains a significant area of interest and active investigation for both the scientific community and medical practitioners. Such advancements are crucial as they serve to expand the available arsenal of tools and techniques for targeted neuromodulation and broaden the spectrum of their potential capabilities in addressing a wider range of neurological and psychiatric conditions. In essence, it is unlikely that any single NIBS method will prove universally capable of addressing all the diverse needs for treating the complex array of brain disorders. Rather, the collective ensemble of methods, each possessing unique and complementary features, provides a rich spectrum of possibilities for precisely modifying neural function for therapeutic purposes. Within this context of continuous innovation and the synergistic application of diverse NIBS approaches, TRFS, leveraging its key features – namely its bimodal neuromodulatory capability, its potential for reaching deep brain structures, and its large spatial span of influence – offers a set of beneficial capabilities that can significantly contribute to the evolving landscape of non-invasive neuromodulation therapies.

## Methods

### Animals

Adult wild-type C57BL/6JxFVB and PV-cre mice (20-35 gr; B6 PV^cre^, The Jackson Lab 017320) were obtained from the Charles River and the Jackson Laboratories, respectively. Mice were kept in cages in a 12hr regular cycle vivarium room dedicated to mice in up to five-occupancy cages.

In all experiments, each animal served as its own control, no randomization or blinding was employed. No prior experimentation had been carried out on the animals.

Implanted mice were moved to single occupancy cages to avoid both social conflicts between animals and damage to the implanted materials.

All experiments were conducted in accordance with the Institutional Animal Care and Use Committee (IACUC) of New York University Medical Center.

### Mice head-fixation

A 3D-printed mice head-fixation set-up with a matching head-mount cap was used (the design files can be found at the following link https://github.com/omidyaghmazadeh/3D_Print_Designs/tree/main/Mouse_HeadFixation_SetUp). This set-up was used for fiber photometry experiments and temperature measurements. Before starting the experiments, animals were trained to habituate with head-fixation for incremental duration from 5min to up to 2hrs.

### Radio frequency stimulation device

Radio frequency wave at 945MHz was radiated from custom design and in-house fabricated ‘stub’ antennas. These antennas were made by unshielding the tip (∼8mm) of a coaxial cable which is connected to a matching circuit. The matching component is a simple single-layer PCB which contains a 50Ω transmission line and a capacitive trace. The length of the capacitive trace is adjusted for impedance matching (by carefully shaving the metallic trace and avoiding leaving sharp edges). In the second version of the antenna, we incorporated a miniature RF connector (type MMCX) so that it could be plugged/unplugged. When fabricated in this way, the front end of the antenna is small and lightweight and can be attached to animal’s skull for consistent stimulation outcome and allowing experimentation in freely moving mice.

Antennas were fed by an RF signal that was generated by a compact USB controlled signal generator (SynthHD, Windfreak Tech. LLC) and amplified by a narrowband high-power amplifier (ZHL-100W-13+, Mini-circuits). In the case of RF stimulation with an unattached antenna, the antenna is placed on top of the skull in a designated location in animal’s 3D printed headcap (Fig.1B). In the case of attached antenna, the tip of the antenna is cemented to animal’s skull (Fig.1F).

### fMRI experiments

#### The MRI scanner

All MRI experiments were performed on a Biospec 70/30 micro-MRI system (Bruker – Billerica MA, USA) equipped with a zero-helium boil-off 300 mm horizontal bore 7-Tesla (7-T) superconducting magnet (300 MHz) based on ultra-shield refrigerated magnet technology (USR). The magnet is interfaced to an actively shielded gradient coil insert (Bruker BGA-12S-HP; OD=198-mm, ID=114-mm, 660-mT/m gradient strength, 130-μs rise time) and powered by a high-performance gradient amplifier (IECO, Helsinki – Finland) operating at 300A/500V. This installation is controlled by an Avance-3HD console operated under Paravision 6.1 and TopSpin 3.1.

#### Surgical Procedure

A mouse head fixation setup with a matching head-mount cap, was incorporated into the cradle design used for imaging experiments. Mice underwent a brief surgery to attach the head cap to their skulls. For this set of experiment, the headcap also served for holding the tip of the TRFS antenna in place (Suppl. Fig.1A) to ensure consistent stimulation. During the surgery, mice were kept anesthetized under a steady stream of isoflurane (2%). The rectal temperature was kept constant at 36–37 °C with a DC temperature controller (TCAT-2, Physitemp LLC, Clifton, NJ), and stages of anesthesia were maintained by confirming the lack of vibrissae movements and nociceptive reflex. The skin of the head was shaved and cleaned by three successive applications of povidone-iodine surgical scrub solution and alcohol pads. The skin was retracted after a medio-sagittal incision, and the bone surface was cleaned with hydrogen peroxide (2%) and allowed to dry. Metabond (Parkell Inc., Edgewood, NY) was applied to the skull surface, and the custom-designed 3D-printed plastic head cap was then attached to the animal’s skull using dental cement. Metabond is fast to dry, and the applied layer of Metabond was completely dry when the cap was positioned and dental cement was applied. Dental cement must be applied between the edges of the cap and the skull; however, in these experiments, we also covered all exposed areas of the skull. Covering the whole skull area is not mandatory, though, for example, when brain implants are used in the experiment (e.g., for optogenetic stimulation). Animals were allowed to recover from surgery (for at least 4 days) before conducting any experiment.

#### Animal preparation

At the start of the experiment, mice were first anesthetized in an induction box supplied with a mixture of 0.5–2% isoflurane and air. Next the animal was moved to a stereotaxic system for a quick surgery to implant a temperature sensor. First, the protective layer on the previously attached headcap was removed. Then a small craniotomy was made and an optical temperature sensor (PRB-100-01M-STM, Osensa Innovation Corp.) was inserted in the brain (at -0.5 mm AP, -0.5 mm ML, and -0.6 mm DV (with a 20 angle) referenced from the Bregma point; Suppl. Fig. 1B) and secured to the skull by dental cement. Next the animal was removed from the stereotaxic system to be placed inside the imaging cradle. The animal’s head was fixed using the appropriate tooth bar and ear bars to avoid possible head motion during the scan. The nose cone was attached to the isoflurane vaporizer system (EZ-155, World Precision Instruments Inc.) by a single 2 mm-diameter plastic tube and a male Luer connector. Animal’s body temperature and respiration rate were monitored throughout the scan using a rectal temperature probe (SA Instruments) and a pressure-sensitive respiration pad (SA Instruments), respectively. The animal’s body temperature was maintained at a constant level (36° to 38 °C) using a heating pad consisting of tubes filled with hot water, connected to a water heater circulating system (Thermo Fisher). An 86 mm inner diameter birdcage coil (Bruker Corp.) was used in transmit mode in combination with a surface loop coil (ID = 20 mm; Bruker Corp.) for signal acquisition in RX mode. The Rx loop coil was placed on top of the animal’s head as close as possible, but without touching it, and secured in place using surgical tape. The antenna tip was then positioned in place and the temperature probe was connected to the recording system. The cradle was then inserted inside the scanner, using the Bruker Autopac system, for brain imaging.

#### MR data acquisition

The imaging experiments began with a FLASH sequence to verify the animal’s head centering, followed by a morphological T2-weighted Turbo-RARE sequence (with two averages and a turbo-factor of 8) using the following parameters: TE/TR = 35/2516 ms; FOV = 30 x 9 mm²; Matrix size = 256 x 256; No. of Axial Slices: 19; Slice Thickness: 0.8 mm; Planar Resolution: 0.117x0.035 mm^2^;Scan Time: 2 min 41 sec. Afterward, fMRI data were collected. Both resting-state (rs-fMRI) and TRFS stimulation-induced (TRFS-fMRI) data were acquired using T2*-weighted single-shot gradient echo – echo planar imaging (GE-EPI) sequence^36,37^ with the following imaging parameters: TE/TR:13.7/1500 ms; FOV: 30x9 mm^2^; Matrix size: 128x64; No. of Axial Slices: 14; Slice Thickness: 0.8 mm; Planar Resolution:0.234x0.140 mm^2^;Scan Time: 7 min 30 sec.

During a recording session, simultaneous RF stimulation (≥25W at 945 MHz) and fMRI imaging (TRFS-fMRI) was performed. Resting-state (rs-fMRI) blocks (without any RF radiation) were also conducted before and after the TRFS-fMRI period. RF stimulation was applied in 15 trials, each consisting of 10s of RF-ON followed by 81s of RF-OFF. Images during the RF-ON epochs were distorted due to RF interference and were discarded from the analysis. To compare TRFS-induced thermal effects on the BOLD signal, data from a high temperature period (the first 15 scans after RF is turned off) were compared with baseline (the last 15 scans before RF is turned on).

#### Data quality check

Quality of the fMRI datasets was vigorously checked similar to our previously published report^38^. In summary, temporal signal-to-noise ratio (tSNR), framewise displacement (FD), and overall brain activity (z-score) during the entire scanning sessions were computed using Matlab-based scripts.

#### fMRI data processing

Data (pre- and post-) processing was performed following the procedure described in our previous report^38,39^.

### TRFS pristine mode

#### Surgical Procedure

Mice were anesthetized with isoflurane and properly positioned (using earbars) in a stereotaxic system. After skin incision and skull cleaning a custom-made 3D printed plastic head-cap base (Clear04 resin, Formlabs) was glued to the skull using dental cement (C&B Metabond, Parkell Inc.). This head-cap base included a guiding element to host the tip of the stub antenna for RF stimulation (Fig. 2A). Prior to the surgery a fiber tip (for fiber photometry) and an optical temperature probe (PRB-100-01M-STM, Osensa Innovation Corp.) were attached to a Microdrive (made using 3D printed parts) by dental cement. Prior to this attachment, the fiber tip was coated with a mixture of 750nl Silk fibroin (5154, Sigma-Aldrich) and 750nl of AAV5-flex-GCaMP6s virus, following the steps as previously described^40^. After making a craniotomy, the prepared fiber and the temperature probe were implanted in mice brain at (AP 1 mm, ML 0.75 mm, DL 0.65 mm) in adult PV-cre mice (B6.129P2-Pvalb^tm1(cre)Arbr^/J, The Jackson Laboratory – strain#017320). The hole around the sensor was filled with a mixture of wax and mineral oil (1.5-2 to 1 mass ratio). A custom-made 3D printed head-cap top was attached to the head-cap base in order to protect the implants. Animals were allowed to recover from surgery before conducting any experiments.

#### Experimental Procedure

Animals were first habituated to head fixation as described earlier. At the time of the measurement, mice were head-fixed to a 3D-printed headfixation setup. The stub antenna V1 was then positioned making sure that the tip of the antenna is sitting in place properly. Fiber cables for temperature and fiber photometry were then connected to the implanted parts. First, temperature measurement was conducted to determine the proper RF power level for the ongoing experiment preparation. RF stimulation (at 945MHz) was applied by first starting with a higher level (6-20 W) to ramp up the temperature increase and then with a lower level (3-8 W) to ensure a plateau-like increased temperature profile (>2°C) during the stimulation (Fig. 2C-E). Next, simultaneous temperature and fiber photometry recording were started. RF stimulation (at 945MHz) with the power levels determined earlier was then applied in an intermittent manner with >25 trials of 45s RF-ON followed by 81s RF-OFF epochs. At the end of the experiment, the fiber cables were disconnected, the antenna was displaced from the headcap, the animal was removed from the head fixation setup, the headcap was covered by parafilm for protection, and the animal was moved to its home cage. At least one Sham session was also conducted with similar parameters as the stimulation sessions with the exception for the power of the RF signal which was at a very low level (0.05W).

### TRFS-induced behavioral changes in pristine mode

#### Surgical Procedure

Mice were anesthetized with isoflurane and properly positioned (using earbars) in a stereotaxic system. After skin incision and skull cleaning a custom-made 3D printed plastic head-cap base (Clear04 resin, Formlabs) was glued to the skull using dental cement (C&B Metabond, Parkell Inc.). Two tips of the front portion of the connector-including stub antenna were then attached to the right and the left sides of the animal’s skull (for hemisphere-different stimulation) using dental cement (Fig. 3A). Two craniotomies (at 1 mm AP, ±1.5 mm ML coordinates), for later temperature measurements, were made. The craniotomies were then covered by a removable silicone gel (Kwik-Sil, World Precision Instruments). The 3D printed head-cap was wrapped with parafilm for protection. Animals were allowed to recover from surgery before conducting any experiments.

#### Temperature Measurement Procedure

Before executing the behavior tests, temperature measurements were conducted to determine the RF power level required for each antenna to induce (>1°C) hemispheral temperature differences required to induce rotation behavioral changes. Animals were first habituated to headfixation as described earlier. At the time of the measurement, mice were head-fixed to a 3D-printed head fixation setup. Their headcap cover and the silicone gels covering the craniotomies (formerly made during the surgery) were removed. Using stereotaxic holders, two fiber optic temperature sensors (PRB-100-01M-STM, Osensa Innovation Corp.) were then inserted at a depth of -2mm DV. RF energy (945MHz) was then fed into each antenna separately and temperature measurements were conducted for different RF power levels (5-25 W). Appropriate power levels, sufficient to induce adequate differential temperature rises between the two hemispheres, were noted after analysis the recoded data. At the end of the experiment, temperature probes were extracted from the brain, the craniotomies were covered with silicone gel (Kwik-Sil, World Precision Instruments), the animal was removed from the headfixation set-up, the headcap was covered by parafilm for protection, and the animal was moved to it home cage.

#### Experimental Procedure

For the behavioral test, animals were first injected with MK-801 (0.3 mg/Kg Intraperitoneal (IP)) to induce hyperlocomotion. One of the implanted antennas (for the side of interest in that session) was then attached to the RF circuit and the mouse was placed inside the open field arena (Fig. 3F). Video recording was then started. Usually, animals were active and moving around in the first few minutes due to being exposed to a novel environment, and then their activity level reduced. However, sometime between 15-30 minutes after the injection, animals showed an increased level of movement because of the drug.

At this point, RF energy (at 945 MHz) at the power level specific to the antenna (identified by temperature measurements) was applied intermittently: approximately 30 trials of 20s of RF-ON followed by 100s of RF-OFF epochs were conducted. Recording was stopped about an hour later, the RF coaxial cable was disconnected, the headcap was covered by parafilm for protection and the animal was returned to its home cage.

### TRFS RF-genetics mode

#### Surgical Procedure

Mice were anesthetized with isoflurane and properly positioned (using earbars) in a stereotaxic system. After skin incision and skull cleaning a custom-made 3D printed plastic head-cap base (Clear04 resin, Formlabs) was glued to the skull using dental cement (C&B Metabond, Parkell Inc.). The tip of the front portion of the connector-including stub antenna was then attached to the right side of the animal’s skull (for hemisphere-different stimulation) using dental cement. Prior to the surgery a fiber tip (for fiber photometry) and an optical temperature probe (PRB-100-01M-STM Osensa Innovation Corp.) were attached to a Microdrive (made using 3D printed parts) by dental cement. Before this attachment, the fiber tip was coated by a mixture of 1500nl Silk fibroin (5154, Sigma-Aldrich), 800nl of AAV9-SYN-mTRPV1-P2A-mCherry virus, and 700nl of AAV1-syn-GCaMP7f virus following the steps as described in^40^. After making a craniotomy, the prepared fiber and the temperature probe were implanted in the mouse brain at (AP 1 mm, ML 0.75 mm, DL 0.70 mm) in adult wild-type mice (C57BL/6). The hole around the sensor was filled with a mixture of wax and mineral oil (1.5-2 to 1 mass ratio). A custom-made 3D-printed head-cap top was attached to the head-cap base to protect the implants (as illustrated in Fig. 4A). Animals were allowed to recover from surgery before conducting any experiments.

#### Experimental Procedure

Animal were first habituated to head fixation as described earlier. At the time of the measurement, mice were head fixed to a 3D-printed head fixation setup. The tip of the antenna attached to the mouse’s skull was then connected to the main part of the stub antenna V2. Fiber cables for temperature and fiber photometry were then connected to the implanted parts and simultaneous temperature and fiber photometry recording were started. RF stimulation (at 945 MHz) with different power levels (1-15 W, inducing varying temperature rises during stimulation) was then applied intermittently, with 15 trials of 20s RF-ON followed by 100s RF-OFF epochs. At the end of the experiment, the fiber cables and the antenna tip were disconnected, the animal was removed from the headfixation set-up, the headcap was covered by parafilm for protection, and the animal was moved to it home cage. At least one Sham session was also conducted with similar parameters as the stimulation sessions with the exception for the power of the RF signal which was at a very low level (0.05 W).

### TRFS-induced behavioral changes in RF-genetics mode

#### Surgical Procedure

Mice were anesthetized with isoflurane and properly positioned (using earbars) in a stereotaxic system. After skin incision and skull cleaning a custom-made 3D printed plastic head-cap base (Clear04 resin, Formlabs) was glued to the skull using dental cement (C&B Metabond, Parkell Inc.). Two tips of the front portion of the connector-including stub antenna were then attached to the right and the left sides of the animal’s skull (for hemisphere-different stimulation) using dental cement (Fig. 5A). Two craniotomies, both of them located either on the right or the left hemisphere (one at 1mm AP, ±1.5 mm ML and another one at 0 mm AP, ±2 mm ML coordinates) were made. 1200 nl of AAV9-SYN-mTRPV1-P2A-mCherry virus was then injected (using a microinjector) through each craniotomy at the depths of 1.9 mm and 2 mm DV for the 1mm AP and the 0 mm AP ones, respectively. This double injection approach was chosen to ensure TRPV1 overexpression over a larger striatal volume. Another craniotomy was performed to mirror the existing one with 0 mm AP coordinates against the midline (at ±2 mm ML). The two craniotomies at this AP level were later used for temperature measurement. After the injection, all craniotomies were covered by a removable silicone gel (Kwik-Sil, World Precision Instruments). The 3D printed head-cap was wrapped with parafilm for protection. Animals were allowed to recover from surgery before conducting any experiments, and a three-weeks waiting time for expression of viral construct was applied.

#### Temperature Measurement Procedure

Before executing the behavior tests, temperature measurements were conducted to determine the RF power level required for each antenna to induce (>1° C) hemispherical temperature differences required to induce rotation behavioral changes. Animals were first habituated to head-fixation as described earlier. At the time of the measurement, mice were head-fixed to a 3D-printed head fixation setup. Their headcap cover and the silicone gels covering the craniotomies (formerly made during the surgery) were removed. Using stereotaxic holders, two fiber optic temperature sensors (PRB-100-01M-STM, Osensa Innovation Corp.) were then inserted at a depth of 2.2 mm DV. RF energy (945MHz) was then fed into each antenna separately and temperature measurements were conducted for different RF power levels (3-25 W). Appropriate power levels, sufficient to induce adequate differential temperature rises between the two hemispheres, were noted after analysis of the recorded data. At the end of the experiment, temperature probes were extracted from the brain, the craniotomies were covered with silicone gel (Kwik-Sil, World Precision Instruments), the animal was removed from the head-fixation set-up, the headcap was covered by parafilm for protection, and the animal was moved to it home cage.

#### Experimental Procedure

For the behavior test, animals were first injected with MK-801 (0.3 mg/Kg IP) to induce hyperlocomotion. One of the implanted antennas (for the side of interest in that session) was then attached to the RF circuit and the mouse was placed inside the open field arena (Fig. 5E). Video recording was then started. Usually, animals were active and moving around in the first few minutes due to being exposed to a novel environment, then reduced their activity. However, sometime between 15-30 minutes after the injection, animals showed an increased level of movement because of the drug. At this point, RF energy (at 945 MHz) at the power level specific to the antenna, identified in temperature measurements, was applied intermittently: 20 to 30 trials of 20s of RF-ON followed by 100s of RF-OFF epochs were conducted. Recording was terminated about an hour later, the RF coaxial cable was disconnected, the headcap was covered with parafilm for protection, and the animal was returned to its home cage.

### Statistical analyses

Statistical analyses were performed by MATLAB functions or custom-made scripts. No statistical methods were used to pre-determine sample sizes, but our sample sizes are similar to those generally employed in the field. All data presented were obtained from experimental replicates with at least three independent experimental repeats for each assay. Statistical comparisons were performed using nonparametric two-tailed Wilcoxon rank-sum (equivalent to Mann-Whitney U test) or Wilcoxon signed-rank test. Due to experimental design constraints, the experimenter was not blind to the manipulation performed during the experiment.

## Reporting summary

Further information on research design is available in the Nature Portfolio Reporting Summary linked to this article.

## Data availability

The data that support the findings of this study are available from the corresponding author upon reasonable request.

## Code availability

Custom MATLAB (MathWorks) scripts were used for data analysis. They are available from the corresponding author upon reasonable request.

Software package used for extraction of animal position/posture were downloaded from https://github.com/DeepLabCut/DeepLabCut.

## Acknowledgements

This work was supported by the following grants: NIH-R01 (# 1R01NS113782-01A1) and partially by NYU CTSI TL1 postdoctoral fellowship (# 2TL1TR001447-06A1) to O.Y.

## Authors Contributions

O.Y. and G.B. conceived the project. L.A. and O.Y. designed the stub antenna. O.Y., L.A., T.M.A., and J.Z. designed the MRI experiments, O.Y., T.M.A., and Z.B.Y. performed MRI experiments and O.Y. and T.M.A. analyzed the corresponding data. O.Y. designed and performed all other experiments and the corresponding data analysis. O.Y. wrote the first draft of the paper and all authors worked on its revisions.

## Ethics declaration

Competing interests:

Authors declare no competing interest.

Experiments were conducted in accordance with the Institutional Animal Care and Use Committee (IACUC) of New York University Medical Center.

## Additional information

## Supplementary Information - Results

### Magnetic Resonance Imaging (MRI) assessment of the effects of TRFS in vivo

To perform fMRI experiments, we designed a 3D print structure for rodent conditioning for MRI experiments (publicly available at: https://github.com/omidyaghmazadeh/3D_Print_Designs/tree/main/3D_Print_Platform_for_In_Vi <u>vo_MRI_in_Mice_and_Rats</u>)^23^. Wild-type (C57BL/6 strain; n=14) mice were anesthetized by IP injection of urethane (1.5g/kg). A 3D-printed head-cap (designed for RF stimulation using our stub antenna; Suppl. Fig.1A) was attached to their skulls and an optical temperature probe was implanted in their brain (Suppl. Fig.1B). They were then head-fixed to our 3D-print small animal conditioning cradle (Suppl. Fig.1C-D). RF stimulation (950MHz; 20W) was applied in an intermittent manner (15 pulses of 15s ON followed by 81s OFF) so that brain temperature increased by approximately 2°C during the RF ON periods (Suppl. Fig.1E-G). Resting-state and TRFS-induced fMRI data were acquired using T2*-weighted single-shot gradient echo–echo planar imaging (GE-EPI) sequence^24,25^. Imaging data were processed using our pipeline described in a former report^26^ and TRFS-induce BOLD signal changes were evaluated. Suppl. Fig.1F shows the resulting temperature change due to the TRFS-paradigm and Fig. 1G shows the average temperature change over RF stimulation pulses (15s ON 81s OFF, total of 64 scans). Due to interference between the RF stimulus and the scanner’s RF coil, images during the RF-ON periods (10 out of 64 scans) were noisy and not interpretable. Therefore, we compared the BOLD signal between the 15 first scans of each stimulation cycle (total of 225; High Temperature (HT) scans) where the brain temperature was most elevated with the last 15 scans (total of 225; Low Temperature (LT) scans) where the brain temperature has lowered (ΔT(avg)=1.41C; Suppl. Fig.1H-I). Suppl. Fig.1J illustrates the percentage of change in the BOLD signal (BOLD_HT-BOLD_LT) in various axial planes across the brain (n=14 mice). Both positive and negative changes in the BOLD signal in response to the brain temperature increase were observed. This is in line with the results from a previous fMRI study on the evaluation of BOLD signal in response to heating caused by high-power light pulses in an experimental set-up for optogenetic stimulation^27^, where authors report a mixture of both negative and positive responses. However, the dominant negative response was likely due to changes in the temperature-dependent MRI parameters such as T_2_*. The positive BOLD response was likely associated with a combination of neural response and physiological changes (e.g. local blood flow^27^) in the brain. Suppl. Fig.2A-C illustrates changes in the BOLD signal in an example area with positive response in a statistically significant manner.

## Supplementary Figures

**Supplementary Figure 1.**
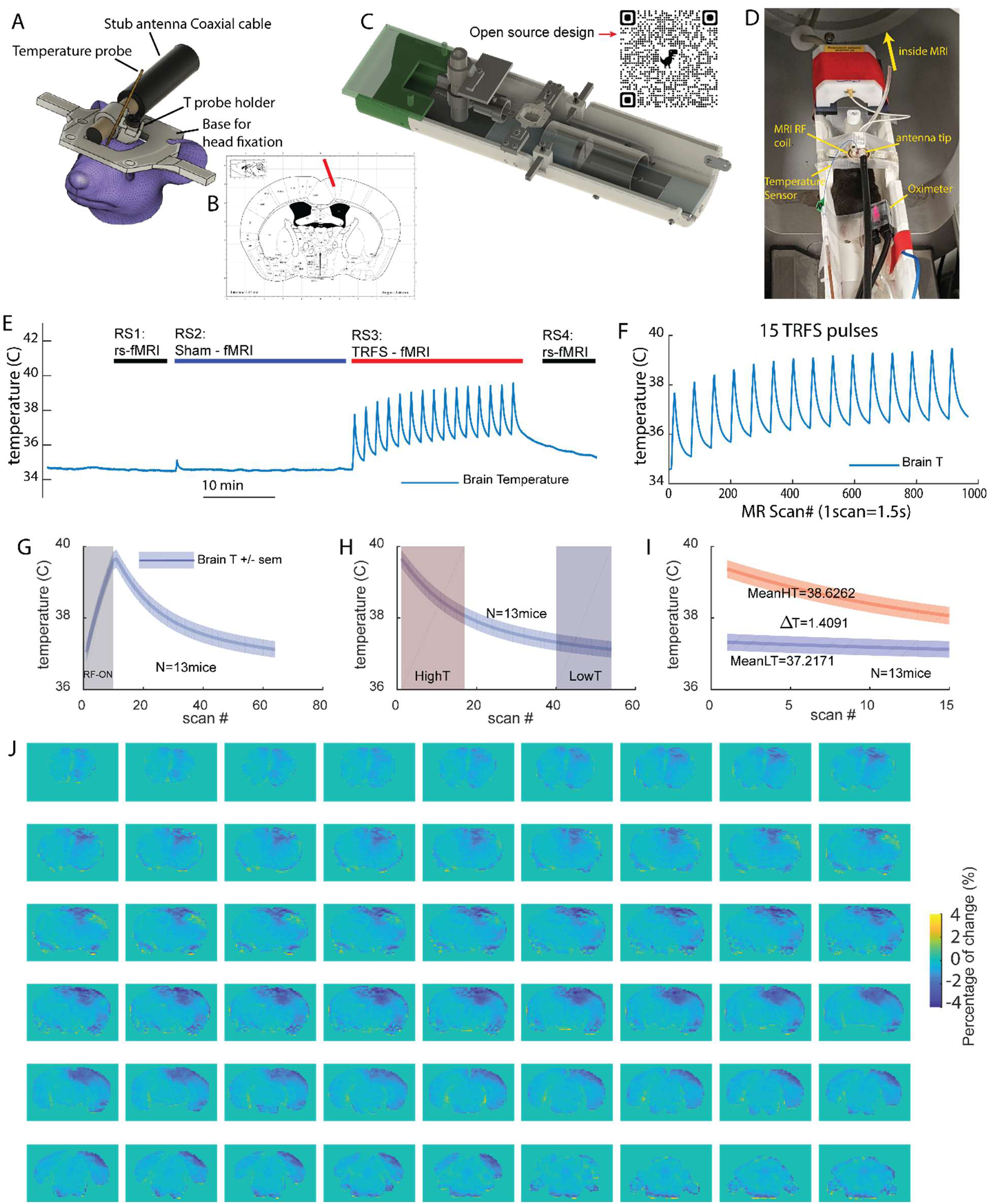
fMRI experiments to evaluate the effect of TRFS o the brain. **A)** Head-cap for placement of antenna tip. **B)** Temperature probe implant coordinates. **C)** 3D print design for animal conditioning. **D)** Photo of the experimental set-up prior to putting the animal in the scanner. **E)** Temperature profile of an example session. **F)** Temperature profile of a 15 pulse TRFS paradigm. **G)** Average temperature profile over an RF pulse (15s ON 81s OFF). H-I) Selecting 15 first (total 225) and 15 last (total of 225) scans in all RF-OFF periods to compare BOLD signal in High Temperature (HT) vs. Low Temperature (LT) periods. **J)** the percentage of change in the BOLD signal (BOLD_HT-BOLD_LT) in various axial planes across the brain (n=14 mice).

**Supplementary Figure 2.**
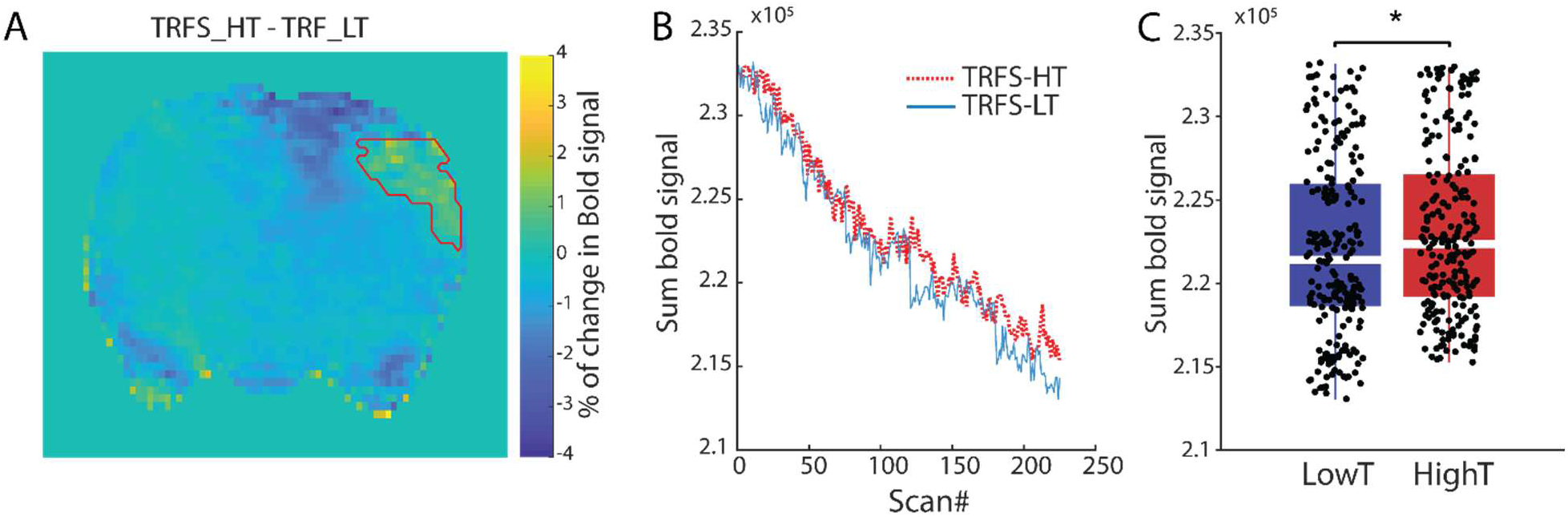
Example plane with positive change in BOLD signal with increasing temperature. **A)** Example brain scan including the example area with positive BOLD response to increasing temperature. **B-C)** Comparison of the BOLD signal for HT and LT scans (data from n=14 mice).

**Supplementary Figure 3.**
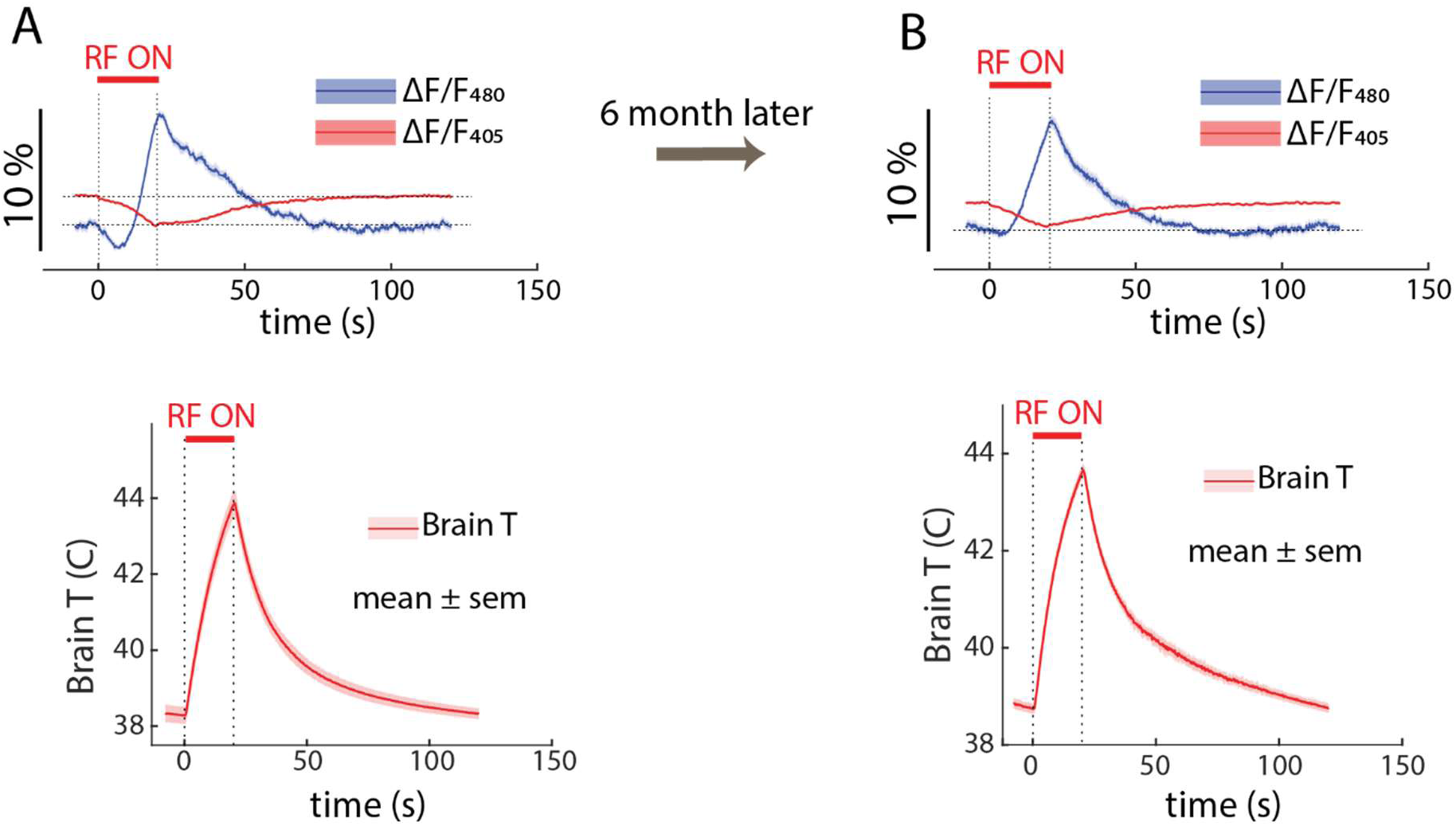
TRFS *RF-genetics mode* effectiveness is preserved at 6month after the initial expression. **A)** top: average fiber-photometry response (mean ± sem) for all (15) trials including the main excitation signal (480 nm, blue) and the isosbestic control signal (405 nm, red) and bottom: the corresponding average (mean ± sem) temperature profile over all the trials measured right after initial expression of TRPV1 channels (∼4weeks after the injection). **B**) Similar to A with data collected 6 months later. Note that the effectiveness of the neuro-excitation is comparable 6 months after the first recordings.

## Supplementary Videos

Supplementary Video 1. **TRFS *pristine mode* induces directional preference ipsilateral to the stimulation side.** Extract of recorded video (played at 4x speed) from animal behavior and TRFS induced directional changes for RF stimulation on the left side. Note the LED light indicating the RF stimulation period when it is ON. Note the preference of the animal to rotate in CCW direction (ipsilateral to the stimulation side) after TRFS is applied.

Supplementary Video 2. **TRFS *pristine mode* induces directional preference ipsilateral to the stimulation side.** Extract of recorded video (played at 4x speed) from animal behavior and TRFS induced directional changes for RF stimulation on the right side. Note the LED light indicating the RF stimulation period when it is ON. Note the preference of the animal to rotate in CW direction (ipsilateral to the stimulation side) after TRFS is applied.

Supplementary Video 3. **TRFS *RF-genetics mode* induces directional preference contralateral to the stimulation side.** Extract of recorded video (played at 4x speed) from animal behavior and TRFS induced directional changes for RF stimulation on the left side which has previously been overexpressed with TRPV1 channel. Note the LED light indicating the RF stimulation period when it is ON. Note the preference of the animal to rotate in CW direction (contralateral to the stimulation side) after TRFS is applied.

Supplementary Video 4. **TRFS *RF-genetics mode* induces directional preference contralateral to the stimulation side.** Extract of recorded video (played at 4x speed) from animal behavior and TRFS induced directional changes for RF stimulation on the right side which has previously been overexpressed with TRPV1 channel. Note the LED light indicating the RF stimulation period when it is ON. Note the preference of the animal to rotate in CCW direction (contralateral to the stimulation side) after TRFS is applied.

